# The Impact of SARS-CoV-2 nsp14 Proofreading on Nucleoside Antiviral Activity: Insights from Genetic and Pharmacological Investigations

**DOI:** 10.1101/2024.07.24.604948

**Authors:** Ju-Yi Peng, Fred Lahser, Christopher Warren, Xi He, Edward Murray, Dai Wang

## Abstract

Nucleoside analogues are a class of well-established antiviral agents that act by being directly incorporated into the viral genome during the replication process, resulting in chain termination or the induction of lethal mutations. While many nucleoside analogues have exhibited broad-spectrum activity against a wide range of viruses, their effectiveness against SARS-CoV-2 is limited. The lack of activity is hypothesized to be attributed to the proofreading function of viral nsp14 exonuclease. In this study, the role of the nsp14 proofreading in modulating nucleoside antiviral activity was investigated using genetic and pharmacological approaches. Introduction of exonuclease attenuation or disabling mutations to nsp14 led to either severe replication defect or increased sensitivity of SARS-CoV-2 and SARS-CoV replicons to specific nucleoside analogues. In contrast, repurposing of HCV NS5A inhibitors to suppress nsp14 exonuclease activity is insufficient to enhance the potency of nucleoside analogues. These findings provided further support for nsp14 as a target for SARS-CoV-2 antiviral development and highlighted the complex interplay between nsp14 proofreading and RNA replication.

## Introduction

Nucleoside analogues (NUCs), which are modified forms of natural nucleosides, have been broadly used for treatment of viral infections, including herpesvirus (1), human immunodeficiency virus (HIV) (2), hepatitis C virus (HCV) (3) and influenza virus (4). These synthetic compounds closely resemble nucleosides or nucleotides in structure. As a result, they can be mistakenly recognized by viral polymerases as a substrate and incorporated into the nascent viral genome during replication (5, 6). The incorporated NUCs may act as chain terminators, preventing further extension of viral genome. Alternatively, they may induce lethal mutations, leading to faulty viral replication and reduced viral fitness (5, 6).

Given the fact that the active sites of viral polymerases, which are responsible for incorporating NUCs into the growing viral genome, are often conserved across different viruses, many NUCs are broadly active against diverse viral species (5, 6). Remdesivir, an adenosine analogue prodrug, was initially developed for the treatment of Ebola virus disease and has been repurposed for treatment of SARS-CoV-2 (7, 8). The broad-spectrum activity of NUCs have made them an essential component in pandemic preparedness and response (5, 9).

Among the licensed NUC drugs, remdesivir, ribavirin, sofosbuvir, tenofovir and favipiravir were predicted by molecular docking to have inhibitory activity against SARS-CoV-2 RNA dependent RNA polymerase (RdRp) (10). Subsequent studies using polymerase extension assays demonstrated that those compounds can be incorporated by SARS-CoV-2 RdRp (11, 12), disrupting the RNA synthesis by direct chain termination (11) or accumulation of deleterious mutations in viral genome (12). However, in the cell-based assays, inhibition of SARS-CoV-2 replication by those compounds were low or nondetectable (13–15).

The limited activity of the NUCs against SARS-CoV-2 has been postulated to be a result of proofreading functions provided by viral encoded exonucleases (5, 6, 16). The presence of exonuclease activity enables viruses to identify and remove misincorporated NUCs from the genome, thereby reducing NUCs intrinsic potencies (6, 17, 18). Coronaviruses are one of few RNA viruses that possess a proofreading function (19, 20). The exonuclease domain resides in the N-terminal region of nonstructrual protein 14 (nsp14) which contains five conserved catalytic residues, D90/E92 (motif I), E191 (motif II), D273 (motif III), and H268 in the active site (21, 22). The exonuclease activity can also be modulated by nsp10, which is associated with nsp14 in the replication complex (19, 20). The proofreading mechanism plays an essential role in maintaining replication fidelity and the stability of the large RNA genome (20, 23), but may also confer natural resistance to NUCs (5). Several recent studies have reported that inhibiting nsp14 exonuclease activity could potentially enhance the antiviral activity of NUCs against coronaviruses (18). Combination of NUCs with HCV NS5A inhibitors, which were predicted to target SARS-CoV-2 nsp14 exonuclease domain, increased the antiviral potency of NUCs against SARS-CoV-2 (13, 14).

While biochemical and cell-based in vitro studies yielded key insights into the impact of nsp14 proofreading function on NUCs antiviral potency (12, 14, 24, 25), these findings also need to be further confirmed through genetic approaches. Phenotypic analysis of specific exonuclease mutations in context of viral replication offers a more direct and precise investigation of these effects. It has been shown that, in murine hepatitis virus (MHV) , MERS-CoV, and SARS-CoV, mutations impairing the exonuclease activity was able to render these viruses more susceptible to inhibition by ribavirin and 5-fluorouracil compared to their wild-type parents (18, 23, 26, 27). However, the genetic evidence supporting the role of proofreading in modulating the NUCs activity against SARS-CoV-2 is currently lacking.

In this study, a panel of mutant SARS-CoV-2 and SARS-CoV replicons with reduced or abolished exonuclease activity was constructed. Alanine substitutions of conserved residues within the active site of the exonuclease are lethal to SARS-CoV-2 replication, while mutations of residues on the nsp14/nsp10 interface resulted in a varying degree of susceptibility or increased sensitivity to NUCs, such as ribavirin and sofosbuvir, depending on the specific mutations. However, inhibition of the SARS-CoV-2 proofreading function with HCV NS5A inhibitors failed to enhance the antiviral potency of NUCs. While our study suggested that combination of nsp14 and nsp12 inhibitors holds promise for treating coronavirus infections, it also argued that in order to fully leverage the benefits of combination therapy, the development of a novel and potent nsp14-specific compound is necessary.

## Materials and Methods

### Molecular docking procedure

The 2D chemical structures for HCV NS5A inhibitors were obtained from PubChem (https://pubchem.ncbi.nlm.nih.gov/) (Fig. S1). All subsequent steps were performed within Schrödinger Maestro software package. Ligprep was used to generate 3D structures of the ligands and to expand their tautomeric and ionic states using the OPLS4 force field at pH 7.0 ± 2.0. Receptor grids were prepared from three high-resolution crystal structures of the SARS-CoV-2 nsp14 enzyme with Protein Data Bank (PDB) codes: 7qgi (nsp14 alone), 7qif (nsp14 alone) and 7diy (nsp14+nsp10). All waters, ions and native ligands were removed and the proteins were prepared by filling in missing side chains with PRIME, performing H-bond assignment using PROPKA at pH 7.0, and running a restrained minimization. Receptor grids were defined via a 10 Å^3^ inner box (30 Å^3^ outer box) centered on active site residues D90, E92, E191, D273, and H268 using the OPLS_2005 force field with no additional constraints. Glide docking was performed in standard-precision (SP) mode with flexible ligand conformational sampling. Post docking PRIME MM-GBSA binding energy calculations were performed on top scoring poses using VSGB solvation model and OPLS4 force field. Glide gscore and MM-GBSA dG Bind energies are reported for top scoring poses.

### Cell lines

Human lung carcinoma cell line A549 (ATCC), human lung epidermoid carcinoma cell line Calu-1 (ATCC), and human hepatoma cell line Huh-7 (ATCC) were cultured in high-glucose Dulbecco’s modified Eagle’s medium (DMEM) (Invitrogen). All cell culture media were supplemented with 10% fetal bovine serum and 1% penicillin/streptomycin. The cells were cultured at 37 °C with 5% CO_2_.

### Construction of SARS-CoV replicon

The replicon sequence is derived from the SARS-CoV isolate (GenBank: AY278741). SARS-CoV replicon were constructed as previously described (5). To construct the bacmid, five fragments (F1 to F5) were synthesized and cloned into a high-copy-number pUC57-kan plasmid by Genewiz (South Plainfield, NJ). These fragments spanned the T7 promoter, SARS-CoV-EGFP-Luc, polyA/RbZ/T7 terminator regions, and included 30-60 base pair overlapping sequences with homology and restriction enzyme sites (Table S1). Three-step cloning was performed to construct the SARS-CoV replicon. pSMART BAC vector (Lucigen) was first linearized by NotI digestion. F1 and F5 fragments were released from pUC57-Kan through MluI digestion. The three fragments were then ligated together using the NEBuilder HiFi DNA Assembly Kit (NEB). The resulting mixture was transformed into BAC-optimized replicator v2.0 electrocompetent cells (Lucigen), yielding pSMART BAC F (1,5). Next, pSMART BAC F (1,5) was linearized with AatII and AscI. F2 and F4 fragments were cloned into pSMART BAC F (1,5) using the same method, resulting in pSMART BAC F (1,2,4,5). Finally, F3 digested with SwaI was cloned into pSMART BAC F (1,2,4,5), resulting in the creation of a full-length non-infectious SARS-CoV replicon.

### Construction of SARS-CoV-2 and SARS-CoV nsp14 mutant replicons

To introduce nsp14 mutations into SARS-CoV-2 Wuhan or SARS-CoV replicons, BAC recombineering was performed as previously described (28, 29). First, an ampicillin (Amp) cassette fragment was PCR amplified by using primers SARS-1-NSP14-AMP-AscI-F/SARS-1-NSP14-AMP-AscI-R or SARS-2-NSP14-AMP-AscI-F/SARS-2-NSP14-AMP-AscI-R with template pcDNA 3.1 (+) vectors (Table S2). Next, parental replicon was transformed into SW102 *E. coli* host along with an Amp fragment, resulting in the creation of nsp14-Amp-replicon. Finally, a fragment containing the nsp14 mutation sequence was synthesized with a 30 bp overlap with the upstream and downstream regions of the nsp14 sequence by Genewiz (South Plainfield, NJ). Following AscI (NEB) digestion, the nsp14-Amp-replicon DNA was ligated to the fragment containing the nsp14 mutations using Gibson Assembly Cloning Kit (NEB). This resulted in the generation of nsp14 mutants-replicons.

### Compound testing

Replicons RNA were prepared as previously described (28). A total of 1× 10^6^ cells were electroporated with 2 µg of replicon RNA and subsequently plated into 96-well plates (Corning; 3904) with a seeding density of 1x10^4^ cells/well. Immediately after electroporation, a 4-fold serial dilution of compounds was added to the wells with DMSO serving as the solvent control. At 48 hours post-transfection, GFP signals were measured by Acumen eX3 scanner. EC_50_ was determined by nonlinear four-parameter curve fitting using GraphPad.

### FRET-based biochemical assay

To investigate the inhibition of exonuclease activity of nsp14 mutant proteins by HCV NS5A inhibitors, the FRET-based biochemical assay was performed as reported previously with modifications (30). Nsp14 wild-type (wt) and mutant proteins, nsp10 proteins and ssRNA oligos were synthesized by GenScript (Piscataway, NJ). To prepare the double-stranded RNA (dsRNA) substrates, RNA oligos (5’6FAM-UCUUUUCGGCCCA-3’; 5’-AAAUAGGGCCGAAAAGA-3’BHQ1) were annealed at a final concentration of 25 μM. The annealing process involved heating the samples in a PCR cycler to 95 °C for 5 minutes, followed by a gradual cooling to 5 °C with 5 °C decrements over 18 cycles of 1-minute each. Exonuclease activity assays were performed at 37 °C in black bottom 96 well plates. The reactions were carried out in a buffer containing 50 mM TRIS-HCl pH 7.5, 2 mM MgCl_2_, and 2 mM DTT. Nsp14 and nsp10 were used at 100 nM and 300 nM, respectively, maintaining a molar ratio of 1:3. The dsRNA substrate was added at a final concentration of 1 μM. The fluorescence intensity of each well was measured every 300 s on the microplate reader (Tecan M1000 Pro) over the course of the activity assay with excitation at 490 nm (± 9 nm) and emission at 520 nm (± 20 nm).

### Quantification and Data Analysis

Data processing and statistical analysis were performed using Prism version 9.4 (GraphPad Software, California). Results are presented as mean ± standard deviation (SD). Statistically significant differences between groups were determined using the one-way analysis of variance (ANOVA) test with Tukey’s *post hoc* analysis.

## Results

### Phenotype of SARS-CoV-2 or SARS-CoV nsp14 mutant replicons

To investigate the role of proofreading in conferring resistance to NUCs, nine SARS-CoV-2 nsp14 mutant replicons were first created. The mutations are located within either the conserved DEED motifs (D90A, E92A, D90A/E92A, E191A, E191Q, and D273A) (17, 23) or the nsp14/nsp10 interface (K9A, K61A, K139A) (25) (Fig. 1& S2). These residues were also selected based on previous reports that these specific mutations either partially or completely inactivated the nsp14 exonuclease activity in other coronavirus models (17, 23, 25, 27). As shown in Fig. 2a, replicons carrying mutations in the nsp14/nsp10 interface demonstrated impaired but viable replication. In contrast, mutations in the DEED motifs led to a significant replication defect.

**Fig. 1.**
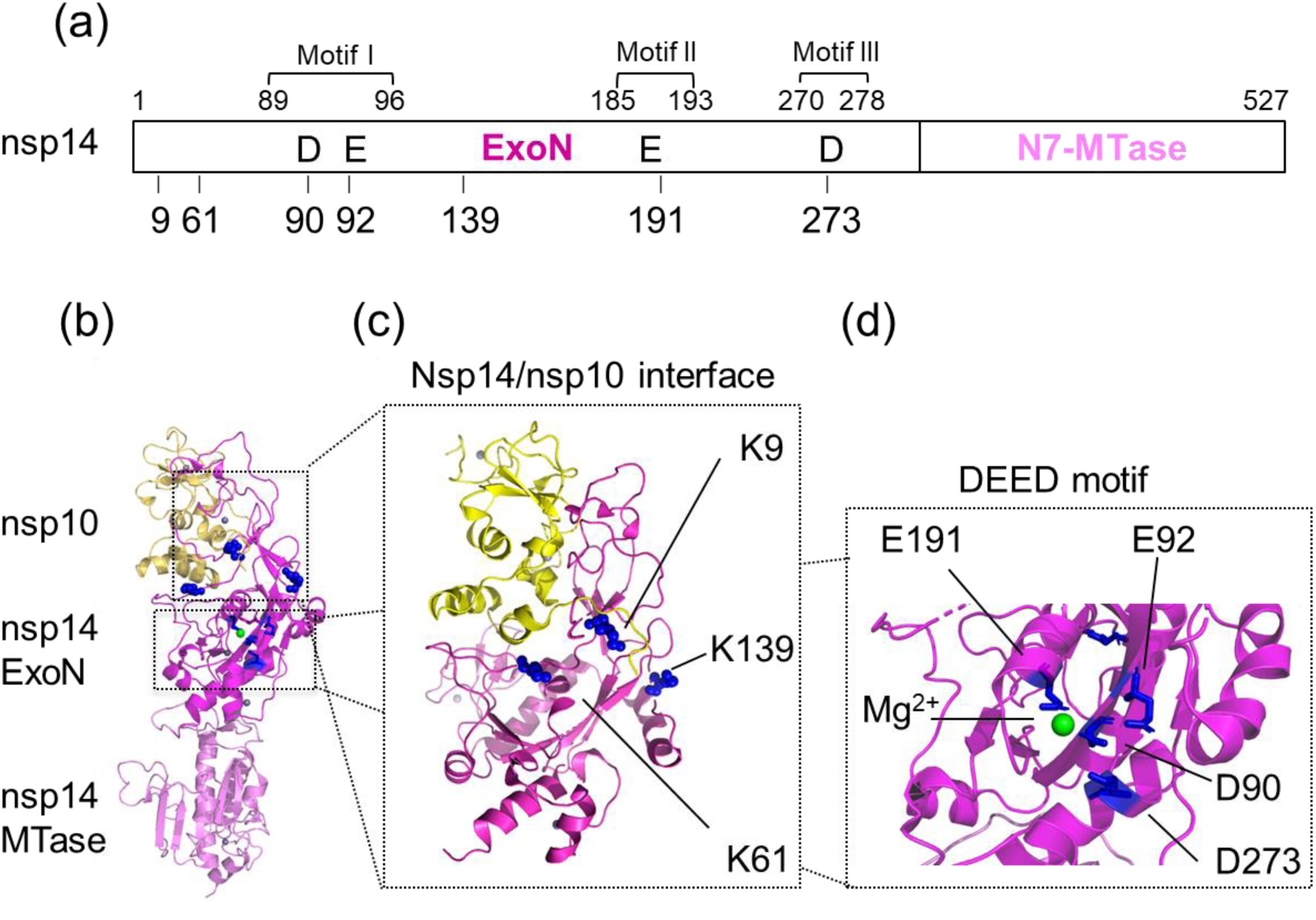
Structure and domain feature of nsp14/nsp10 complex (a) The domains of SARS-CoV-2 nsp14 protein with the position of mutations in this study. (b) 3D structure of SARS-CoV-2 nsp14 and its domains. The exonuclease domain of nsp14 is shown in magenta, MTase domain of nsp14 is shown in light pink, and nsp10 is shown in yellow. (c) Location of the basic patch and the nsp14/nsp10 RNA complex model. Residues K9 (blue ribbon) and K61 (blue ribbon) interact with the N terminus of nsp10, respectively. K139 (blue ribbon) is located farther down along the basic patch. (d) Exonuclease active site in the presence of Mg^2+^ (green ball). Residues D90, E92, E191, and D273 (blue stick) were identified as active-site residues of nsp14 exonuclease.

**Fig. 2.**
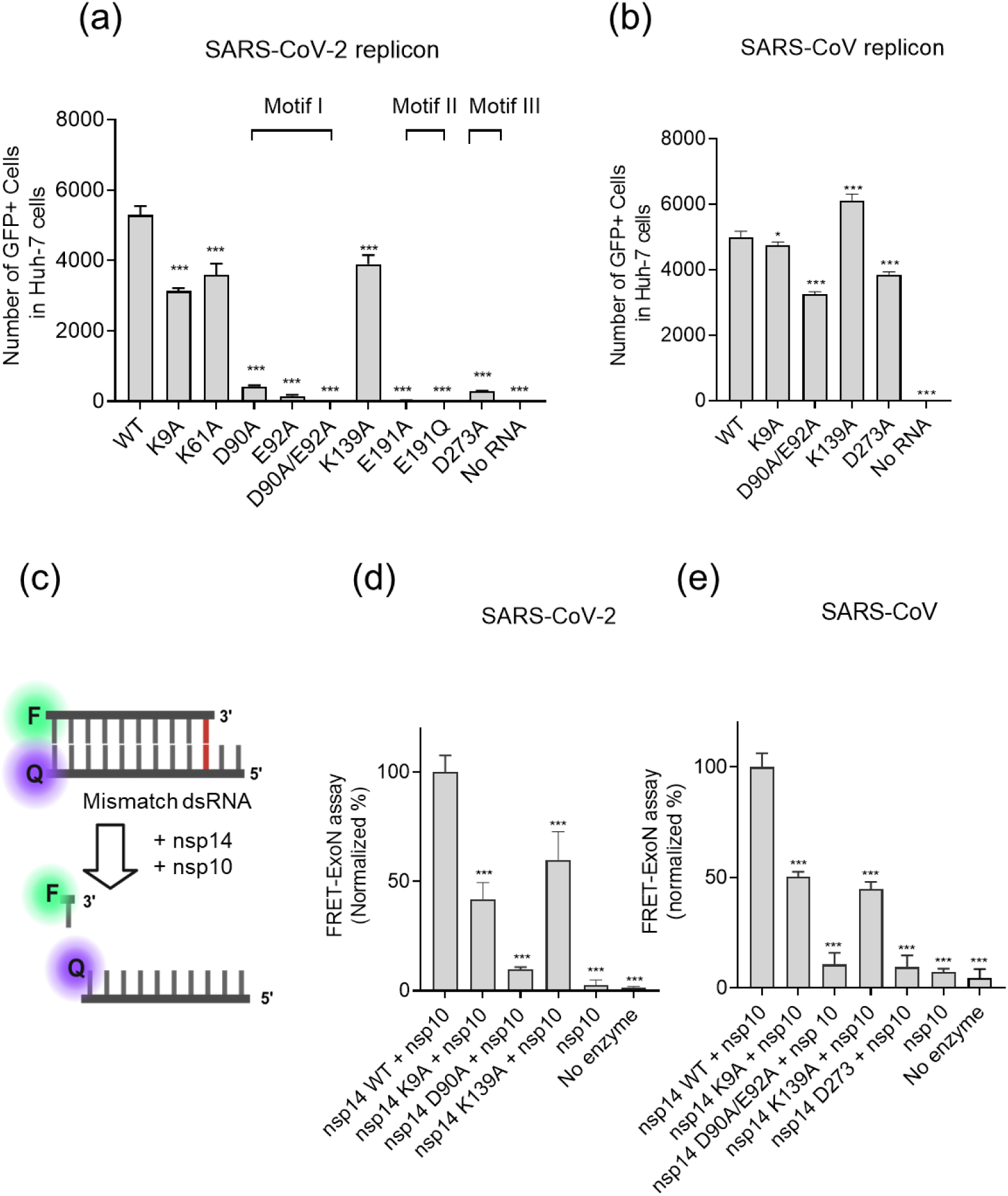
The growth phenotype of nsp14 mutant replicons and the exonuclease activity of corresponding recombinant nsp14 mutant proteins analyzed by the FRET-based biochemical assay. (a & b) Growth phenotype of SARS-CoV-2 and SARS-CoV nsp14 mutant replicons in Huh-7 cells. (c) Schematic illustration of the FRET-based biochemical assay. (d & e) Exonuclease activity of recombinant SARS-CoV-2 and SARS-CoV nsp14 mutant proteins by the FRET-based biochemical assay. Statistical significance of the difference compared with WT and each nsp14 mutation. *, P < 0.05; ***, P < 0.001. The data are presented as mean ± SD.

The viable mutations identified in SARS-CoV-2, such as K9A and K139A, were then introduced to the SARS-CoV replicon. Additional SARS-CoV replicons containing the nsp14 D90A/E92A or D273A mutations were also constructed, since those mutations were viable in SARS-CoV from the previous report (23). NSP14 of SARS-CoV-2 sharing high amino acid similarity with SARS-CoV, and all the substitutions are conserved in these residues. (Fig. S2). As shown in Fig. 2b, all 4 mutant SARS-CoV replicons replicated to high levels compared to the parental replicon. Even though the D90A/E92A or D273A mutations were lethal to SARS-CoV-2 growth (Fig. 2a), they only caused a slight reduction in SARS-CoV replication (Fig. 2b).

### Exonuclease activity of recombinant nsp14 mutant proteins

Next, the viable SARS-CoV-2 and SARS-CoV nsp14 mutants were subjected to the FRET-based biochemical assay to assess the impact of these mutations on exonuclease activity (Fig. 2c). The dsRNA probe, labeled with a fluorophore and a quencher in close proximity, was intentionally designed with the last nucleotide at 3” end mismatched. As the nsp14/nsp10 exonuclease complex recognizes the terminal mismatch in the substrate strand, it progressively excises bases from the 3ʹ end until the two RNA strands separate, resulting in an increase in fluorescence signal. As shown in Fig. 2d, the K9A and K139A mutations in SARS-CoV-2 nsp14 reduced its exonuclease activity by approximately 54% to 56% compared to nsp14 WT. D90A mutant was almost completely deficient in exercising the mismatched nucleoside. A similar trend was also observed in SARS-CoV nsp14 mutants (Fig. 2e). The biochemical data confirmed that the defect of exonuclease activity of the mutants in comparison to nsp14 WT.

### Sensitivity of nsp14 mutant replicons to NUCs

If nsp14 prevents or removes the incorporation of NUCs into the viral genome, the nsp14 knockdown or deficient mutants would be expected to be more sensitivity to NUCs. To test the hypothesis, the potency of NUCs, including remdesivir, ribavirin, sofosbuvir, tenofovir, and favipiravir against the mutant replicons were determined. Those NUCs can be categorized as either chain terminators or mutagenic agents. Sofosbuvir and tenofovir, which are approved for treatment of HCV and HIV infections, respectively, are known as immediate terminators. They lack the 3’-OH group required for the formation of the phosphodiester bond with the next nucleotide, resulting in premature termination of the DNA or RNA chain and inhibition of further chain elongation (5). On the other hand, remdesivir functions as a delayed chain terminator, allowing further extension of the RNA chain after initial incorporation. This compound possesses a modified 1’-cyano group, creating steric hindrance and altering the conformation of the growing nucleic acid chain, ultimately leading to premature termination of RNA synthesis (31). Favipiravir, approved for treating influenza virus, exhibits a dual mechanism of action as both a chain terminator and a mutagenic agent (32, 33). Ribavirin primarily acts as a mutagen and is commonly used in the treatment of HCV and respiratory syncytial virus (RSV) infections (34, 35).

As shown in Fig. 3, SARS-CoV-2 or SARS-CoV nsp14 mutant replicons showed a varying degree of susceptibility or increased sensitivity to the NUCs depending on the specific mutations. Remdesivir was the most potent in inhibiting the growth for both SARS-CoV-2 and SARS-CoV replicons. Mutations introduced to nsp14 however did not significantly alter replicon sensitivity of the remdesivir. Ribavirin also demonstrated activities against both WT replicons. SARS-CoV-2 was more susceptible to ribavirin inhibition compared to SARS-CoV. The nsp14 D273A mutation of SARS-CoV showed the most sensitivity increase to ribavirin. In contrast to remdesivir and ribavirin against SARS-CoV-2, no other compounds were able to inhibit the WT replicon growth by more than 50%. Favipiravir blocked the replicon RNA synthesis only at high concentrations. Limited potency shift was observed against nsp14 mutants. Tenofovir and sofosbuvir were largely ineffective against WT replicons. Mutations in nsp14 did not confer susceptibility to tenofovir, however, the nsp14 D90A of SARS-CoV-2 and D273A of SARS-CoV mutant replicons can be partially inhibited by sofosbuvir at concentrations above 625 nM.

**Fig. 3.**
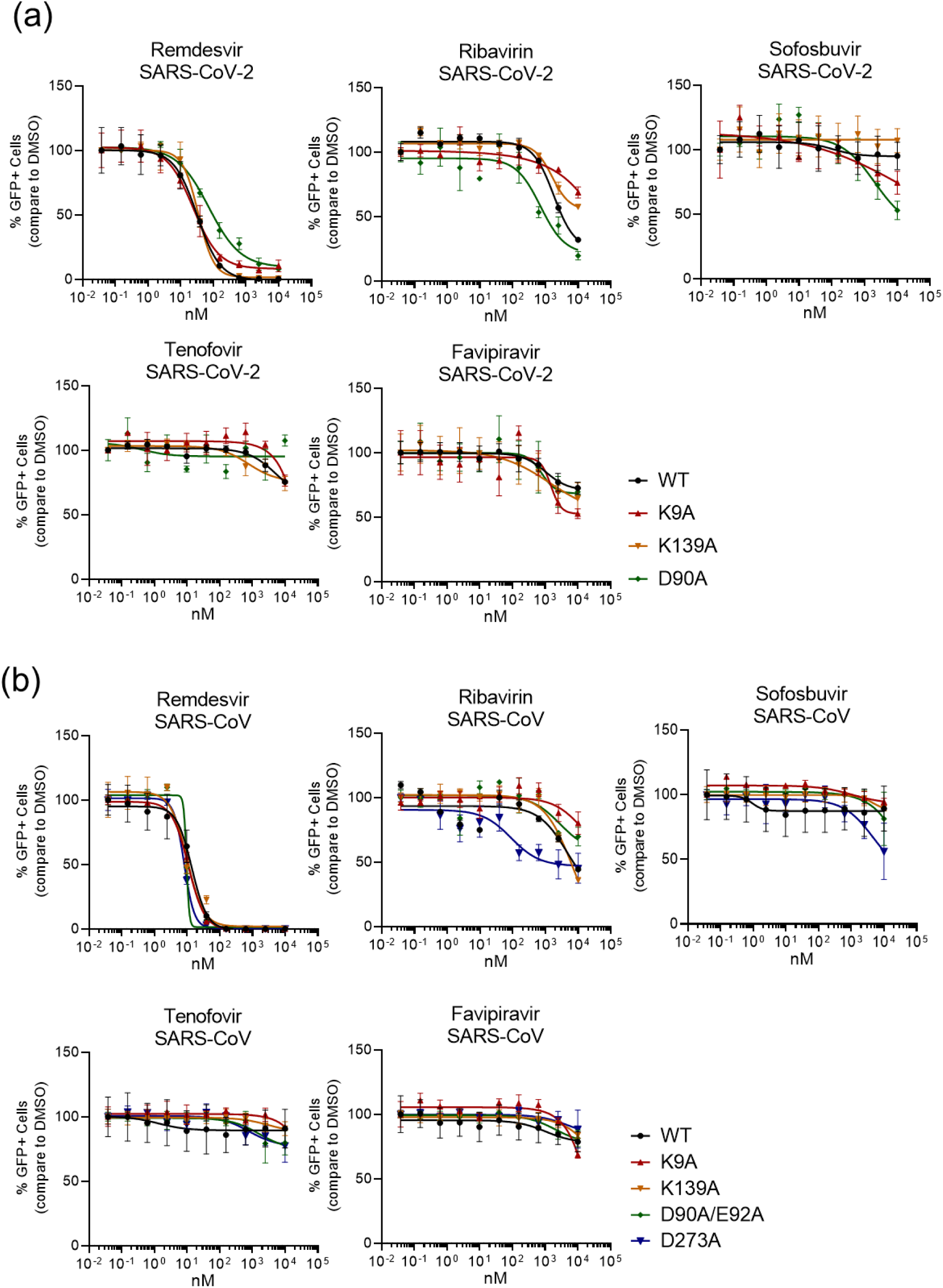
Inhibition of SARS-CoV-2 or SARS-CoV nsp14 mutant replicons by NUCs. Dose-dependent responses of SARS-CoV-2 (a) and SARS-CoV (b) nsp14 mutant replicons reporters’ activity to NUCs in Huh-7 cells. The Huh-7 cells were electroporated with SARS-CoV-2 or SARS-CoV replicon RNA and incubated with compounds at indicated concentrations. The activities were determined by percentage of GFP signals comparing to DMSO-treated cells. The data was shown mean ± SD.

### Inhibition of nsp14 exonuclease activity by HCV NS5A inhibitors

To gain further insights into the relationship between exonuclease activity and NUCs resistance, we next sought to inactivate the nsp14 proofreading function pharmacologically. Currently, the best-studied compounds against SARS-CoV-2 nsp14 exonuclease activity are the HCV NS5A inhibitors (14, 25). Here, the potential of FDA approved HCV NS5A inhibitors, including daclatasvir, velpatasvir, ruzasvir, elbasvir, ombitasvir and pibrentasvir, to inhibit SARS-CoV-2 nsp14 exonuclease was first assessed using computational ligand docking and the FRET-based biochemical assay. Molecular docking of NS5A inhibitors was performed in Glide software using three available high-resolution nsp14 protein structures (Fig. 4a) (36, 37). The docking ability between compounds and proteins were assessed using Glide gscore and MM-GBSA free energy estimates.

**Fig. 4.**
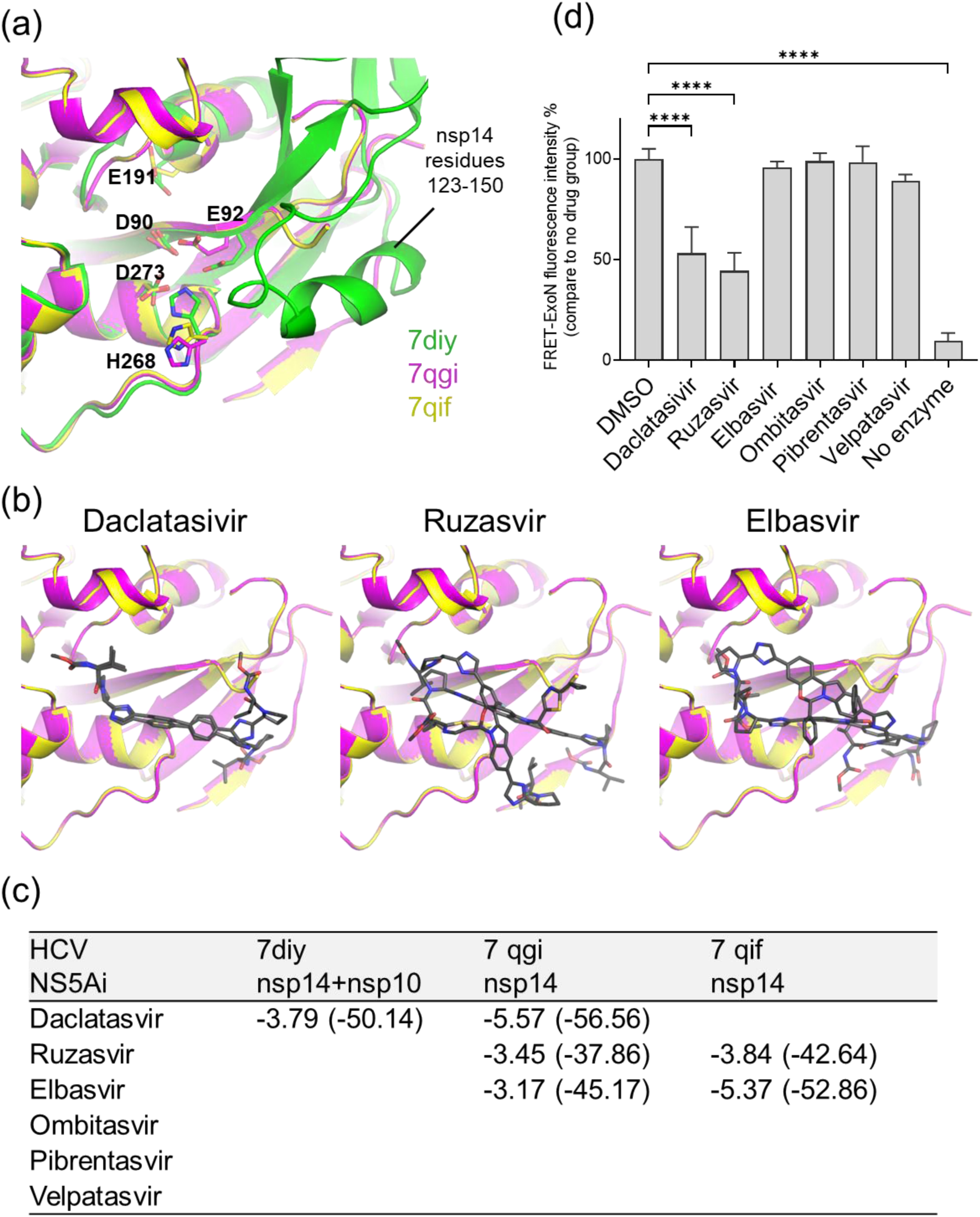
Inhibitory effects of HCV NS5A inhibitors on SARS-CoV-2 nsp14 exonuclease activity. (a) Overlay of active sites from three structural models of nsp14 used for *in silico* docking: 7diy from SARS-CoV-2 nsp10 bound to nsp14 exonuclease domain (green), 7qgi from SARS-CoV-2 nsp14 (magenta), and 7qif from SARS-CoV-2 nsp14 complexed with 7MeGpppG (yellow). Key active site residues are highlighted along with residues 123-150 of nsp14 which become ordered upon nsp10 binding (b) Overlay of the best scoring docking poses of HCV NS5A inhibitors bound to apo nsp14 (7qgi and 7qif) exonuclease active site. (c) Glide gscores and PRIME MM-GBSA dG Bind scores of top poses. (d) Exonuclease activity of the nsp14 in the presence of 10 µM HCV NS5A compounds. DMSO was used as a positive control, and no enzyme served as a negative control. Fluorescence was measured at reaction time of 1500 seconds. The shown data represents a single independent measurement conducted with triplicates. Error bars represent SD.

As shown in Fig. 4b and 4c, the docking procedure yielded a modestly scored, but consistent low energy pose between apo nsp14 (PDB 7qgi and 7qif) and daclatasvir. The ordering of nsp14 residues 123-150 upon binding to nsp10 (Fig. 4a) is predicted to sterically interfere with this pose, leading to poorer scores for daclatasivir binding to the nsp14+nsp10 complex (PDB 7diy) (Fig. 4c). Poses of ruzasvir and elbasvir bound to apo nsp14 were less consistent between structures and yielded worse scores compared to daclatasvir (Fig. 4b and 4c). No low energy poses were identified for velpatasvir, ombitasvir or pibrentasvir. These results indicate that daclatasivir, along with possibly ruzasvir and elbasvir, may act as inhibitors by binding to the active site of the apo nsp14 enzyme and may prevent nsp10-induced active site rearrangement. However, the lack of an available known nsp14 inhibitor structure makes interpretation of these docking results challenging.

The compounds identified by the molecular docking were further tested experimentally. Recombinant nsp14/nsp10 complex was subjected to the FRET-based biochemical assay with or without HCV NS5A inhibitors. Daclatasvir and ruzasvir were capable of inhibiting nsp14 exonuclease activity albeit modestly (47-51% reduction at 10 μM concentration) (Fig. 4d). No activity was observed for pibrentasvir, ombitasvir, velpatasvir or elbasvir against nsp14.

### Lack of inhibitory effect of HCV NS5A inhibitors on SARS-CoV-2 replicon

Next, the potency of the HCV NS5A inhibitors against SARS-CoV-2 replication was determined in the cell-based replicon assay (Table 1). Among the HCV NS5A inhibitors tested, only ombitasvir demonstrated inhibition effect on SARS-CoV-2 replicon growth, although it did not show significant activity at a concentration of 10 μM in the FRET-based biochemical assay (Fig. 4d). The EC_50_ values of ombitasvir were found to be 554.4 ± 342 nM, 1155.4 ± 786 nM and 1196.7 ± 585 nM in Huh-7 cells, A549 cells and Calu-1 cells, respectively. In contrast, daclatasvir and ruzasvir, which demonstrated inhibition of nsp14 exonuclease activity in the FRET-based biochemical assay, failed to inhibit the replication of SARS-CoV-2 in the cell-based replicon assay. Given the low potency of daclatasvir and ruzasvir in the FRET-based biochemical assay, it is possible that partially reduction of nsp14 exonuclease is not sufficient to limit SARS-CoV-2 replication.

**Table 1.** Inhibitory effect of HCV NS5A inhibitors on SARS-CoV-2 replicons in different cell lines.

| Compound | Huh-7<br>EC<sub>50</sub> nM | A549<br>EC<sub>50</sub> nM | Calu-1<br>EC<sub>50</sub> nM |
| --- | --- | --- | --- |
| Daclatasvir | >10,000 | >10,000 | >10,000 |
| Ruzasvir | >10,000 | >10,000 | >10,000 |
| Elbasvir | >10,000 | >10,000 | >10,000 |
| Ombitasvir | 554.4 ± 342 | 1155.4 ± 786 | 1196.7 ± 585 |
| Pibrentasvir | >10,000 | >10,000 | >10,000 |
| Velpatasvir | >10,000 | >10,000 | >10,000 |

HCV NS5A: hepatitis C virus's nonstructural protein 5A

### Impact of HCV NS5A inhibitors on NUCs potency against SARS-CoV-2

In light of the inhibition of nsp14 exonuclease activity by daclatasvir and ruzasvir biochemically, we next investigated whether these inhibitors were able to enhance the sensitivity of the SARS-CoV-2 replicon to NUCs, similar to the observations from the genetic analysis described above. At a concentration of 10 μM HCV NS5A inhibitors in combination with various concentrations of NUCs, ombitasvir increased remdesivir potency by 1.3 fold, while other combinations presented no synergistic effect (Fig. 5 and Table 2). It remains unknown whether a more potent nsp14 exonuclease inhibitor will render SARS-CoV-2 more susceptible to NUCs.

**Fig. 5.**
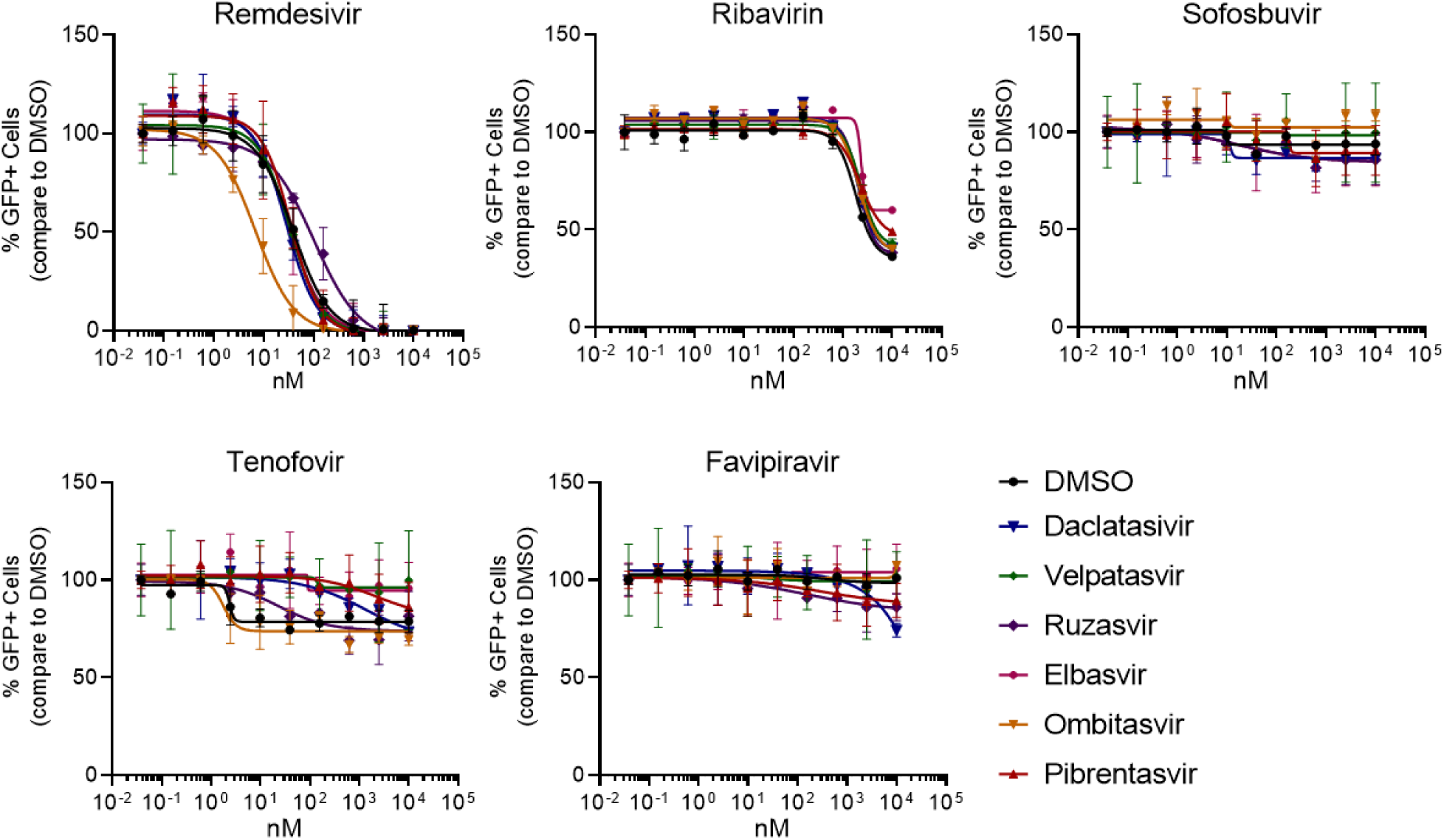
Dose-dependent responses of SARS-CoV-2 replicon reporter activity to NUCs in combination with 10 µM HCV NS5A inhibitors. Huh-7 cells were electroporated SARS-CoV-2 replicon RNA and incubated with compounds at indicated concentrations. The activities were determined by percentage of GFP signals comparing to DMSO-treated cells. The data was shown mean ± SD.

**Table 2.** Antiviral activity of combinations of NUCs with 10 µM HCV NS5A inhibitors on SARS-CoV-2 replicon expression in Huh-7 cells.

| NUCs | HCV NS5Ai | Huh-7<br>EC<sub>50</sub> nM | A549<br>EC<sub>50</sub> nM | Calu-1<br>EC<sub>50</sub> nM |
| --- | --- | --- | --- | --- |
| Remdesivir | DMSO | 38.6 ± 12.2 | 64.2 ± 22.1 | 38.6 ± 13.4 |
| Remdesivir | Daclatasvir | 28.2 ± 6.3 | 72.8 ± 24.6 | 38.7 ± 15.2 |
| Remdesivir | Ruzasvir | 102.1 ± 31.8 | ND | ND |
| Remdesivir | Elbasvir | 30.0 ± 7.3 | ND | ND |
| Remdesivir | Ombitasvir | 6.8 ± 1.5 | 82.1 ± 14.2 | 48.9 ± 25.9 |
| Remdesivir | Pibrentasvir | 35.2 ± 10.8 | 95.7 ± 32.3 | 64.2 ± 23.2 |
| Remdesivir | Velpatasvir | 34.4 ± 10.8 | ND | ND |
| Ribavirin | DMSO | 1820.0 ± 303.6 | ND | ND |
| Ribavirin | Daclatasvir | 2294.0 ± 223.3 | ND | ND |
| Ribavirin | Ruzasvir | 2112.1 ± 188.3 | ND | ND |
| Ribavirin | Elbasvir | 2322.9 ± 241.5 | ND | ND |
| Ribavirin | Ombitasvir | 2091.2 ± 223.2 | ND | ND |
| Ribavirin | Pibrentasvir | 2222.6 ± 230.0 | ND | ND |
| Ribavirin | Velpatasvir | 2002.4 ± 188.2 | ND | ND |
| Sofosbuvir | DMSO | >10,000 | >10,000 | ND |
| Sofosbuvir | Daclatasvir | >10,000 | >10,000 | ND |
| Sofosbuvir | Ruzasvir | >10,000 | ND | ND |
| Sofosbuvir | Elbasvir | >10,000 | ND | ND |
| Sofosbuvir | Ombitasvir | >10,000 | >10,000 | ND |
| Sofosbuvir | Pibrentasvir | >10,000 | >10,000 | ND |
| Sofosbuvir | Velpatasvir | >10,000 | ND | ND |
| Tenofovir | DMSO | >10,000 | >10,000 | ND |
| Tenofovir | Daclatasvir | >10,000 | >10,000 | ND |
| Tenofovir | Ruzasvir | >10,000 | ND | ND |
| Tenofovir | Elbasvir | >10,000 | ND | ND |
| Tenofovir | Ombitasvir | >10,000 | >10,000 | ND |
| Tenofovir | Pibrentasvir | >10,000 | >10,000 | ND |
| Tenofovir | Velpatasvir | >10,000 | ND | ND |
| Favipiravir | DMSO | >10,000 | >10,000 | ND |
| Favipiravir | Daclatasvir | >10,000 | >10,000 | ND |
| Favipiravir | Ruzasvir | >10,000 | ND | ND |
| Favipiravir | Elbasvir | >10,000 | ND | ND |
| Favipiravir | Ombitasvir | >10,000 | >10,000 | ND |
| Favipiravir | Pibrentasvir | >10,000 | >10,000 | ND |
| Favipiravir | Velpatasvir | >10,000 | ND | ND |

ND: not determined

## Discussion

There remains a need for the development of new treatment options for SARS-CoV-2. While several antivirals, such as remdesivir, molnupiravir, nirmatrelvir/ritonavir have received FDA approval for treatment (38), the emergence of new variants and the risk of viral resistance underscores the importance of continuing efforts for identifying new antiviral therapies. Historically, RdRp and Mpro were prioritized in antiviral development due to their essential functions in SARS-CoV-2 replication (39, 40). However, recent research has also highlighted the potential of nsp14 exonuclease as a druggable target (21, 24, 30). Nsp14 is a component of replicase complex that provides proofreading function ensuring the accuracy of viral replication (19, 20). Inhibition of the exonuclease activity has been proposed as a strategy to introduce errors in the viral genome, which may reduce viral fitness and replication (19, 23, 27). Alternatively, inhibition of the exonuclease activity of nsp14 also enhances the incorporation of NUCs into viral genome, increasing the susceptibility of the virus to the antiviral effects of NUCs (19, 23, 27).

Both SARS-CoV-2 and SARS-CoV nsp14 proteins contain the DEED motifs in their exonuclease domains (20, 21). These motifs are conserved and required for the catalytic activity of the enzyme (20, 21). Alanine substitution of the DEED residues resulted in near-complete inactivation of exonuclease activity in a FRET-based biochemical assay. SARS-CoV viruses (23) or replicons (Fig. 2b) containing those mutations are generally viable but growth impaired. To our surprise, the same mutations were found to be lethal to SARS-CoV-2 replicon (Fig. 2a), indicating the proofreading activity of SARS-CoV-2 might play a more critical role in maintaining the stability of the viral genome compared to SARS-CoV, and the potential of the DEED motifs as a target for developing antivirals against SARS-CoV-2. In contrast to the DEED mutants, SARS-CoV-2 and SARS-CoV replicons with alanine substitution of the residues involved in nsp14/nsp10 interaction showed a partially reduction of exonuclease activity and attenuated replication but remained viable.

Only a limited impact of the exonuclease mutations on the sensitivity of SARS-CoV-2 and SARS-CoV to NUCs was observed (Fig. 3). The extent of this impact is dependent on the specific virus, mutation, and compound being studied. For SARS-CoV-2, K139A and D90A mutants showed a slight increase of sensitivity to sofosbuvir. No potency enhancement was observed for remdesivir, ribavirin, tenofovir and favipiravir (Fig. 3a). For SARS-CoV, the most notable shift in drug potency was detected in nsp14 D273A replicon treated with ribavirin (Fig. 3b). It is still unclear why only D273A mutations enhanced viral sensitivity to ribavirin despite of a similar level of impaired exonuclease activity was observed in D90A/E92A mutant (Fig. 3b). Both exonuclease activity and the interaction between the replicase complex and compounds may play a role in determine the NUCs potencies.

To gain further insight into the mechanisms underlying the shift in sensitivity to NUCs, we explored the pharmacological inhibition of nsp14 exonuclease activity. However, the availability of specific small molecule compounds targeting nsp14 exonuclease activity is currently limited. Previous molecular docking analyses have identified HCV NS5A inhibitors, such as pibrentasvir, ombitasvir and daclatasvir, that have the potential to bind the active site of SARS-CoV-2 nsp14 and interfere with its exonuclease activity (14, 41). In those studies, the predicted 3D structure of SARS-CoV-2 nsp14 inferred from its ortholog in SARS-CoV was used as its definitive structure has not been solved when the analyses were conducted (14, 41). Only daclatasvir and ruzasvir are confirmed to have inhibitory effects against nsp14 determined by the FRET-based biochemical assay. Here, we re-analyze the interactions between HCV NS5A inhibitors and SARS-CoV-2 based on recently solved, high-resolution structures of nsp14. Among the HCV NS5A inhibitors tested in this study, daclatasvir, ruzasvir, and elbasvir demonstrated modest docking scores to the apo nsp14 models, whereas velpatasvir, pibrentasvir and ombitasvir are not predicted to bind nsp14.

In the cell-based replicon assays, however, neither daclatasvir nor ruzasvir showed inhibitory effect on SARS-CoV-2 replication. In contrast, ombitasvir which did not inhibit the exonuclease activity in the FRET-based biochemical assay reduced the viral replication in Huh-7, A549 or Calu-1 cells. (Fig. 4 and Table 1). Ombitasvir might have a multi target effect on SARS-CoV-2 as it has also been predicted to interact with Mpro and Spike protein (42, 43). Partial inhibition of nsp14 exonuclease activity by daclatasvir and ruzasvir did not boost the sensitivity of SARS-CoV-2 against NUCs tested either. This finding, together with the mutagenesis data (Fig. 4 & 5), suggested that inhibition of exonuclease activity alone might not be sufficient to enhance the NUCs activity. An effective compound may also need to interact with replicase in a specific manner in order to be able to influence the proofreading of NUCs.

A major limitation of this study is that the cell-based replicon assays only allow evaluation of the single-cycle replicon of the virus. As a result, low-level inhibition may not be detectable using this approach and multiple cycles of viral replication are required for the antiviral effect to become apparent. Therefore, it is important to confirm the impact of exonuclease mutations and compounds on viral replication through live virus infections.

In summary, the study provided evidence supporting nsp14 exonuclease as a target for developing antiviral therapies against SARS-CoV-2, either as monotherapy or in combination with NUCs treatments. The investigation revealed that mutations in conserved residues within the catalytic site of nsp14 can be lethal to SARS-CoV-2 replication. Additionally, mutations impacting the interaction between nsp14 and nsp10 demonstrated the potential to enhance the potency of NUCs. The study also suggested that inhibition the exonuclease function alone may not sufficiently lower SARS-CoV-2’s resistance to NUCs, highlighting the challenges of dual-targeting the exonuclease and RdRp.

## Acknowledgements

The authors would like to thank Merck Sharp & Dohme LLC, a subsidiary of Merck & Co., Inc., Rahway, NJ, USA, and the MRL Postdoctoral Research Fellowship Program for providing funding for this work.

