## Supplemental files for "The Impact of SARS-CoV-2 nsp14 Proofreading on Nucleoside Antiviral Activity: Insights from Genetic and Pharmacological Investigations"

Nucleoside analogue

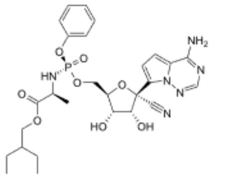

Remdesivir

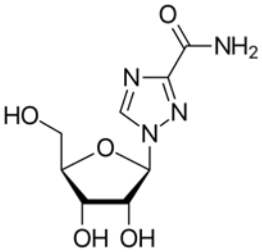

Ribavirin

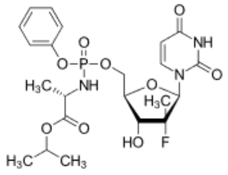

Sofosbuvir

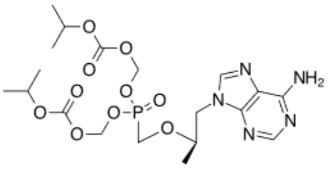

Tenofovir

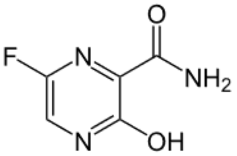

Favipiravir

HCV NS5A inhibitor

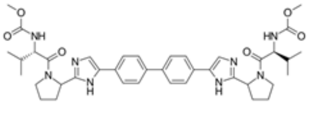

Daclatasvir

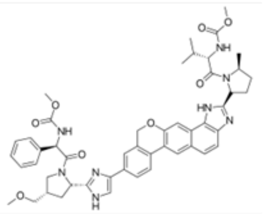

Velpatasvir

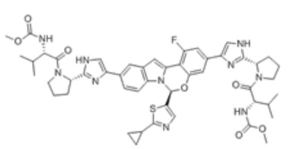

Ruzasvir

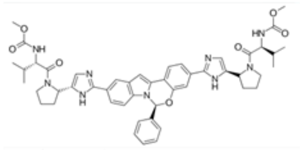

Elbasvir

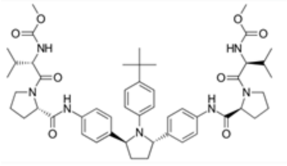

Ombitasvir

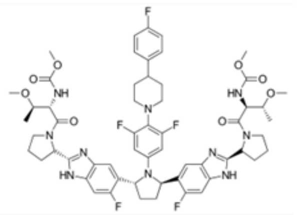

Pibrentasvir

**Fig. S1** Chemical structure of compounds used in this study.

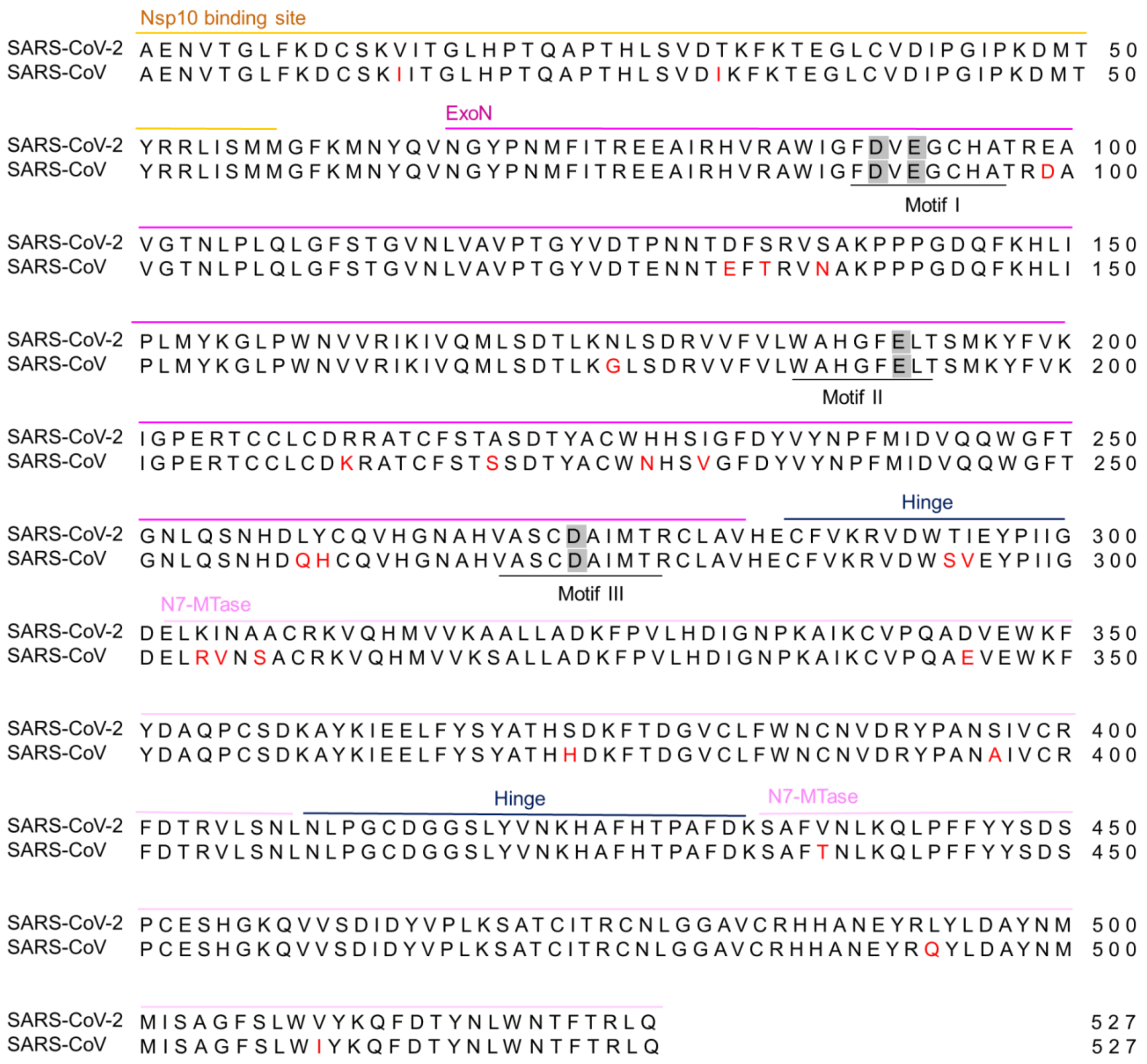


**Fig. S2** Alignment of nsp14 protein sequences of SARS-CoV-2 (NC_045512) and SARS-CoV (AY278741). DEED motifs are shaded in grey color. Non-conserved residues are shown in red (above 95% conservation). The alignment was generated using Clustal Omega.

**Table S1** Sequences of the synthetic cDNA fragments used to construct SARS-CoV replicon

| Fragment | Sequence |
|---|---|
| F1 | 5'- CGCGTTTGACCATGTTGGTATGATTTAAATTCAGTGCGGCCGCTAATACGACTCACTATAGATATTAGGTTTTTACCTACC - 3' |
| | 3'-AGCAAGGTACATTCTTATGTGCGAATGAGTGACGTCATCGGCGCGCCGGACGAGCACTTCTTCATCGTGGACCGGCTACGCGT - 5' |
| F2 | 5'- ACGCGTAGCAAGGTACATTCTTATGTGCGAATGAGTACACTGGTAACTATCAGTGTGGTCATTA -3' |
| | 3'-TTGTTTACAAAGTCTACTATGGTAATGCTTGACGTCATCGGCGCGCCCATAGTGTATACGGCATGCTCTCATGCAGCACGCGT - 5' |
| F3 | 5'- ATTTAAATTTGTTTACAAAGTCTACTATGGTAATGCTTTAGATCAAGCTATTTCCATG - 3' |
| | 3'- CTCTATTACCCATCTGCTCGCATAGTGTATACGGCATGCTCTCATGCAGCATTTAAAT - 5' |
| F4 | 5'- ACGCGTCATAGTGTATACGGCATGCTCTCATGCAGCTGTTGATGCCCTATGTGAAA - 3' |
| | 3'-GACATCGCCTACTGGGACGAGGACGAGCACTTCTTCATCGTGGACCGGCTACGCGT - 5' |
| F5 | 5'- ACGCGTGGACGAGCACTTCTTCATCGTGGACCGGCTGAAGAGCCTGATCAAATACA - 3' |
| | 3'-TAGCTTCTTAGGAGAATGACAAAAAAAAAAAAAAAAAAAAAAAAAAGGGTCGGCATGGCATCTCCACCTCCTCGCGGTCCGACCTGGGCATCCGAAGGAGGACGCACGTCCACTCGGATGGCTAAGGGAGAGCCTGCAGTAGCATAACCCCTTGGGGCCTCTAAACGGGTCTTGAGGGGTTTTTTGGCGGCCGCTTCTATAGTGTCACCTAAATACTAGTGACTCCAGCACGCGT - 5' |

Green: T7 promoter; red: overlapping sequences; blue: HDV ribozyme; purple: T7 terminator; underline: restriction sites

**Table S2** Primer list used to amplify Amp cassette for the nsp14-Amp-replicon

| Primer | Sequence |
| --- | --- |
| SARS-NSP14-AMP-AscI-F | 5'- AGAGATCTTTATGACAAACTGCAATTTACAAGTCTAGAAATACCACGTCGCAATGTGGCTACATTAAAGGCGCGCCGGAACCCCTATTTGTTTATT - 3' |
| SARS-NSP14-AMP-AscI-R | 5'- AGGTGCTTCGCCGGCGTGTCCATCAAAGTGTCCTTTATTAACAACATTATAAGCCACATTTTCTAAACTGGCGCGCCTTACCAATGCTTAATCAGTG - 3' |
| SARS-2-NSP14-AMP-AscI-F | 5'- ATGACAAGTTGCAATTTACAAGTCTTGAAATTCCACGTAGGAATGTGGCAACTTTACAAGGCGCGCCGGAACCCCTATTTGTTTATT - 3' |
| SARS-2-NSP14-AMP-AscI-R | 5'- ACCCTGTTGTCCATCAAAGTGTCCCTTATTTACAACATTAAAAGCCACATTTTCTAAACTGGCGCGCCTTACCAATGCTTAATCAGTG - 3' |

Amp: ampicillin; underline: restriction sites
